# Splice Variants of the SWR1-type Nucleosome Remodeling Factor Domino Have Distinct Functions during *Drosophila melanogaster* Oogenesis

**DOI:** 10.1101/052746

**Authors:** Kenneth Boerner, Peter B. Becker

## Abstract

SWR1-type nucleosome remodeling factors replace histone H2A by variants to endow chromatin locally with specialized functionality. In *Drosophila melanogaster* a single H2A variant, H2A.V, combines functions of mammalian H2A.Z and H2A.X in transcription regulation and DNA damage response. A major role in H2A.V incorporation for the only SWR1-like enzyme in flies, Domino, is assumed, but not well documented *in vivo*. It is also unclear, whether the two alternatively spliced isoforms, *dom-A* and *dom-B*, have redundant or specialized functions. Loss of both DOM isoforms compromises oogenesis causing female sterility. Therefore, we systematically explored roles of the two DOM isoforms during oogenesis using a cell type-specific knockdown approach. Despite their ubiquitous expression, DOM-A and DOM-B have non-redundant functions in germline and soma for egg chamber formation. We show that chromatin incorporation of H2A.V in germline and somatic cells depends on DOM-B, while incorporation in endoreplicating germline nurse cells is independent of DOM. In contrast, DOM-A promotes the removal of H2A.V from stage 5 nurse cells. Remarkably, the two DOM isoforms have distinct functions in cell type-specific development and H2A.V exchange.

**Summary Statement**

Isoforms of nucleosome remodeling factor Domino change chromatin structure by histone variant exchange to direct essential cellular processes in oocyte development.

## Introduction

The local replacement of nucleosomal histone H2A by variants is an evolutionary conserved principle that endows chromatin with structural and functional diversity (Bönisch and Hake, 2012; Talbert and Henikoff, 2010). Among the different H2A variants two can be considered ‘universal’ since they are found in all eukaryotes: H2A.Z and H2A.X. Currently, H2A.X is best known for the role of its phos-phorylated form in DNA damage signaling. H2A.Z, on the other hand, most likely has a role in regulating transcription through chromatin as it specifically marks the nucleosomes next to nucleosome-depleted regions of promoters. Curiously, *Drosophila melanogaster* only has a single H2A variant, H2A.V, which combines the functions of H2A.Z and H2A.X as architectural element downstream of active promoters, in heterochromatin organization and as phosphorylated form (γH2A.V) in the DNA damage response (Baldi and Becker, 2013).

H2A variants are incorporated by Swi2 ATPase domain related 1 complex (SWR1)-like remodeling enzymes, which use the energy freed by ATP-hydrolysis to disrupt canonical nucleosomes. These remodeling enzymes are evolutionary conserved from yeast to humans. Mammals utilize two different SWR1-like enzymes, E1A-binding protein p400 (p400) and Snf2-related CBP activator protein (SRCAP) (Cai et al., 2005; Eissenberg et al., 2005; Jin et al., 2005; Ruhl et al., 2006). In contrast, the *D. melanogaster* genome only encodes a single SWR1-like enzyme, the *domino* (*dom*) gene (Ruhf et al., 2001). Alternative splicing of the *dom* transcript gives rise to two isoforms, *dom-A* and *dom-B* (Ruhf et al., 2001), which differ in their C-terminal regions. The longer DOM-A isoform features C-terminal poly-glutamine stretches and a SWI3–ADA2–N-CoR–TFIIIB (SANT) domain, whereas the DOM-B C-terminus is largely unstructured (Eissenberg et al., 2005; Ruhf et al., 2001). Early reports found DOM-B rather ubiquitously expressed, but DOM-A was detected only in cells of the embryonic nervous system, larval salivary glands and in S2 tissue culture cells (Eissenberg et al., 2005; Messina et al., 2014; Ruhf et al., 2001) suggesting distinct functions.

SWR1-like remodelers typically reside in multi-subunit complexes (Morrison and Shen, 2009), but little is known about DOM-containing complexes. DOM-A has been purified from S2 cells as part of a 16-subunit assembly also containing the acetyltransferase Tat-Interactive Protein 60 (TIP60), apparently combining features of the yeast SWR1 remodeling and Nucleosomal Acetyltransferase of H4 (NuA4) acetyltransferase complexes (Kusch et al., 2004). Likewise, our understanding of the roles of DOM enzymes in H2A.V exchange reactions *in vivo* is anecdotal. DOM has been linked to incorporation of H2A.V at the E2F promotor (Lu et al., 2007). It has also been suggested that H2A.V exchange requires prior acetylation (Kusch et al., 2004; Kusch et al., 2014).

*Domino* was characterized as a gene required for cell proliferation and viability, homeotic gene regulation and Notch signaling (Eissenberg et al., 2005; Ellis et al., 2015; Gause et al., 2006; Kwon et al., 2013; Lu et al., 2007; Ruhf et al., 2001; Sadasivam and Huang, 2016; Walker et al., 2011). Previous genetic analyses did not attempt to resolve distinct functions of the two isoforms, as the known *dom* mutant alleles affect both splice forms. *Dom* is essential for fly development (Ruhf et al., 2001) and *dom* mutants die during pupariation (Braun et al., 1998; Ruhf et al., 2001). Furthermore, oogenesis in adult flies is strongly perturbed causing sterility (Ruhf et al., 2001).

*D. melanogaster* oogenesis provides an excellent opportunity to study self-renewal and differentiation of germline and somatic stem cells (GSC and SSC, respectively) in the context of egg chamber morphogenesis (Hudson and Cooley, 2014; Ting, 2013; Yan et al., 2014). The formation and maturation of eggs starts in the germarium at the most anterior end of an ovariole, a tubular substructure of the ovary. There, somatic terminal filament and cap cells form a niche for 2-3 GSCs. These cells divide asymmetrically to self-renew and to shed a daughter cystoblast in region 1 of the germarium. This cystoblast initiates four mitotic divisions with incomplete cytokinesis to form an interconnected 16-cell cyst. As this cyst travels through region 2 to the posterior end of the germarium one particular cell is determined to become the oocyte. In region 3 the remaining 15 cells transform into polyploid nurse cells by changing their cell cycle program to endoreplication. In parallel, SSCs in region 2 of the germarium produce somatic follicle cells, which encapsulate 16-cell cysts to form individual egg chambers. Oogenesis further runs through 14 stages of egg chamber development to produce a functional egg.

It is not surprising that nucleosome remodeling factors have been found important for oogenesis given the widespread requirement for chromatin plasticity during development (Chioda and Becker, 2010; Ho and Crabtree, 2010; Iovino, 2014). Interestingly, a germline-specific RNA interference (RNAi) screen identified DOM as important for GSC self-renewal and cystoblast differentiation (Yan et al., 2014). DOM function is also essential for somatic follicle cell development, since *dom* loss-of-function alleles show defects in SSC self-renewal (Xi and Xie, 2005). The mechanisms of DOM function in these cell type-specific processes are unclear, but involvement of H2A.V incorporation and turnover has been suggested. H2A.V levels show a modest decrease in *dom* mutant GSC clones in *D. melanogaster* testes (Morillo Prado et al., 2013). During early stages of oogenesis, H2A.V is ubiquitously expressed in somatic and germline cells, but not detectable in mutant germline clones for MORF-related gene on chromosome 15 (MRG15), a DOM-A/TIP60 complex subunit (Joyce et al., 2011). Yet, direct evidence for a role of DOM – and specific roles for each isoform – in H2A.V incorporation during oogenesis is lacking.

We used a cell type-specific RNAi approach to dissect the functions of DOM-A and DOM-B during *D. melanogaster* oogenesis. Our analysis suggests non-redundant requirement for both DOM isoforms in several different cell differentiation programs and for H2A.V exchange.

## Results

### Expression of both DOM splice variants in germline and somatic cells of *D. melanogaster* ovarioles

Domino is ubiquitously expressed in the female germline of *D. melanogaster* ovarioles (Yan et al., 2014), but other studies suggest expression of only one isoform, DOM-B, in soma and germline cells (Ruhf et al., 2001; Xi and Xie, 2005). Due to the lack of specific DOM-A and DOM-B antibodies for immunofluorescence microscopy (IFM) we generated transgenes expressing GFP-3xFLAG-tagged *domino* from a recombined fosmid using the flyfosmid recombineering technique (Ejsmont et al., 2009). This way, we obtained fly lines expressing N-terminally tagged Domino (*GFP-dom*) and C-terminally tagged DOM-A *(dom-A-GFP)* and DOM-B *(dom-B-GFP)* from its chromosomal regulatory context (Fig. 1A). These latter two constructs express one isoform with a tag while the other one is untagged. As a control we replaced the *domino* locus with the GFP-3xFLAG cassette (Δ*dom-GFP*) (Fig. 1A).

**Fig. 1.**
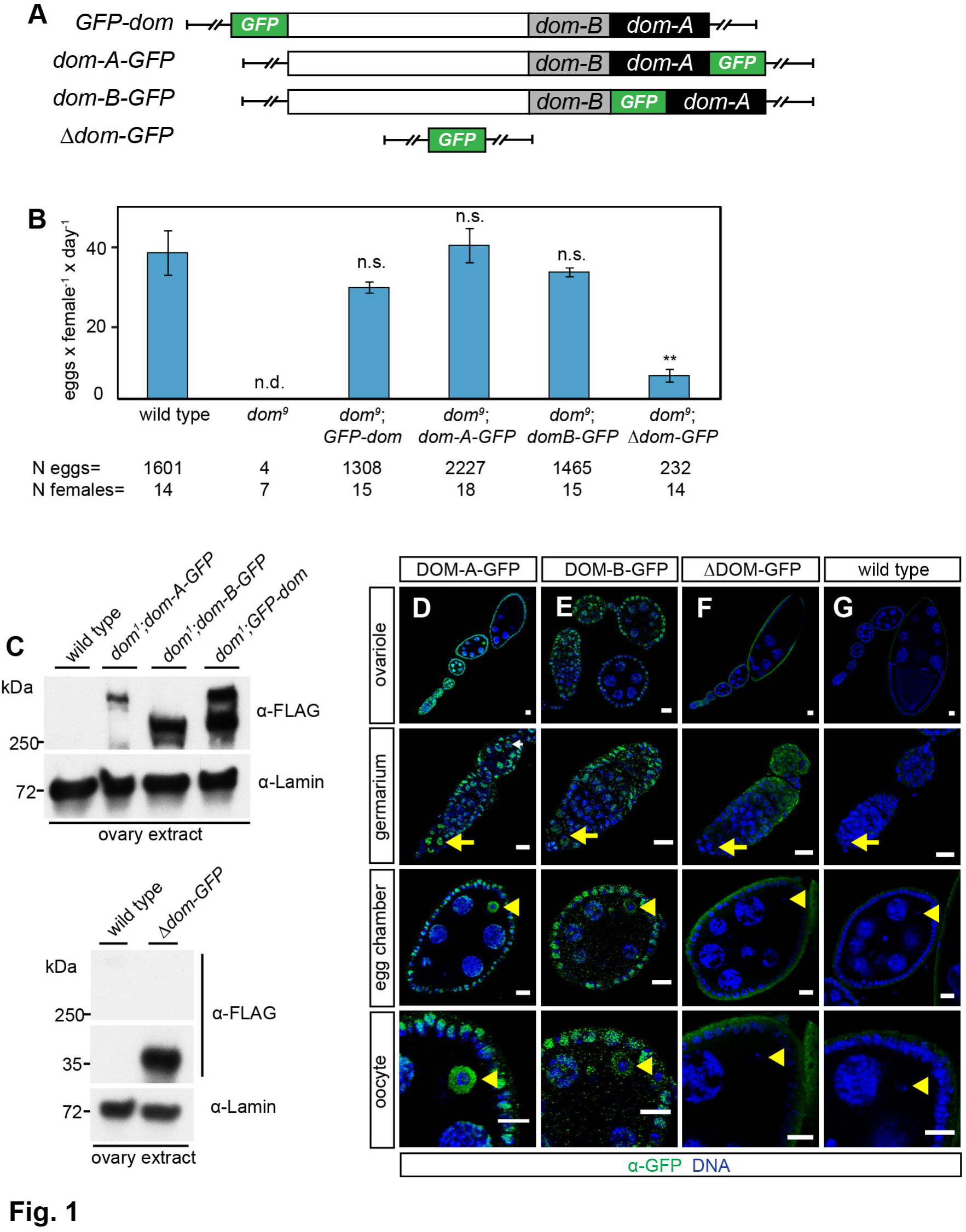
Ubiquitous expression of DOM-A and DOM-B in *D. melanogaster* ovarioles. (A) Schematic representation of the *domino* fosmid constructs. White, black and grey rectangle represent 5’ genomic region of *dom*, splice variant of *dom-A* or *dom-B*, respectively. Green rectangle shows inserted 2xTY1-sGFP-3xFLAG-tag. (B) Complementation of *dom*^*9*^ allele with *dom* transgenes rescues sterility phenotype. Quantification of egg laying capacity is shown. The data show mean values of laid eggs per female and day with SD of three biological replicates. N eggs represents the total number of scored eggs and N females the total number of analyzed females. Two-tailed Student’s t-test was used in comparison to wild type. ** represents a p-value of <0.01, n.s. not significant and n.d. not determined. (C) Western blot from ovaries probed with αα-FLAG is shown for the following homozygous genotypes: wild type, *dom*^*1*^;*dom-A*, *dom*^*1*^;*dom-B*, *dom*^*1*^;*GFP-dom*. Lamin signal served as a loading control. (D-G) Immunofluorescence images of different oogenesis stages with staining of GFP (green) and DNA (blue) are shown for the following homozygous genotypes: (D) *dom-A-GFP*, (E) *dom-B-GFP*, (F) Δ*dom-GFP* fosmid and wild type. Yellow arrow and arrowhead indicate germline stem cells and oocyte nuclei, respectively. Scale bar: 10 μm.

We attempted to complement two different *dom* alleles to assess the functionality of the *dom* transgenes. Loss of Domino in the *dom*^*1*^ allele causes lethality in pupal stage and the *dom*^*9*^ allele is semi-lethal and sterile (Ruhf et al., 2001). Viable homozygous *dom*^*1*^ or *dom*^*9*^ fly lines were obtained by complementation with the *GFP-dom, dom-A-GFP or dom-B-GFP* transgenes (Fig. S1A). In contrast, the Δ*dom-GFP* transgene did not rescue pupal lethality in the *dom*^*1*^ allele and compromised viability in the *dom*^*9*^ allele (Fig. S1A). Furthermore, the characteristic necrotic lymph glands phenotype of *dom*^*1*^ larvae (Braun et al., 1998; Ruhf et al., 2001) was completely rescued by the N and C-terminally tagged *dom* transgenes (Fig. S1B,C). More importantly for this study, egg laying capacity was restored to wild type levels in homozygous *dom*^*9*^ females complemented with the *dom* transgenes, but not by the Δ*dom-GFP* transgene (Fig. 1B). Our analysis provides the first comprehensive validation of *dom* allele phenotypes using complementation with *dom* transgenes.

Western blotting using the FLAG antibody confirmed the expression of *dom* transgenes in ovaries (Fig. 1C) and larval brains (Fig. S1D) (Ruhf et al., 2001). Next, we used the GFP antibody to study the localization of DOM-A and DOM-B in ovarioles. The specificity of antibody was confirmed as wild type ovarioles showed only background staining (Fig. 1G). We confirmed the ubiquitous expression of DOM-B in germline and somatic cells (Fig. 1E) (Ruhf et al., 2001; Xi and Xie, 2005). Interestingly, we also observed expression of DOM-A in germline and somatic cells throughout the entire ovariole (Fig. 1D). Both Domino isoforms were present in GSCs and cystoblasts, SSCs and follicle cells and were enriched in oocyte nuclei in comparison to nurse cells (Fig. 1D,E). Nuclear localization was not due to the GFP-tag since GFP expressed from the Δ*dom-GFP* transgene showed only cytoplasmic signal (Fig. 1F). In summary, both isoforms, DOM-A and DOM-B, are expressed in germline and somatic cells during *D. melanogaster* oogenesis.

### Cell type-specific knockdown of DOM-A and DOM-B reveals requirements during fly development

To dissect the cell type-specific requirements for DOM-A and DOM-B we induced RNAi by expressing small hairpin (sh) RNA directed against *domino* under the control of the Gal4/UAS system (Ni et al., 2011). We also generated transgenic flies expressing *shRNA* directed selectively against each individual DOM isoform. *shRNA* directed against an irrelevant *GFP* sequence served as appropriate control for non-specific effects in all experiments.

Ubiquitous depletion of DOM should resemble the lethal phenotype of the *dom*^*1*^ allele (Ruhf et al., 2001). Indeed, we observed pupal lethality upon expression of *dom-1 shRNA* with an *actin-Gal4* driver, while siblings lacking the driver (*CyO*/*dom-1 shRNA*) were not affected (Fig. 2A). Remarkably, individual depletion of DOM-A or DOM-B led to pupal lethality indicating essential functions for either isoform during fly development (Fig. 2A).

**Fig. 2.**
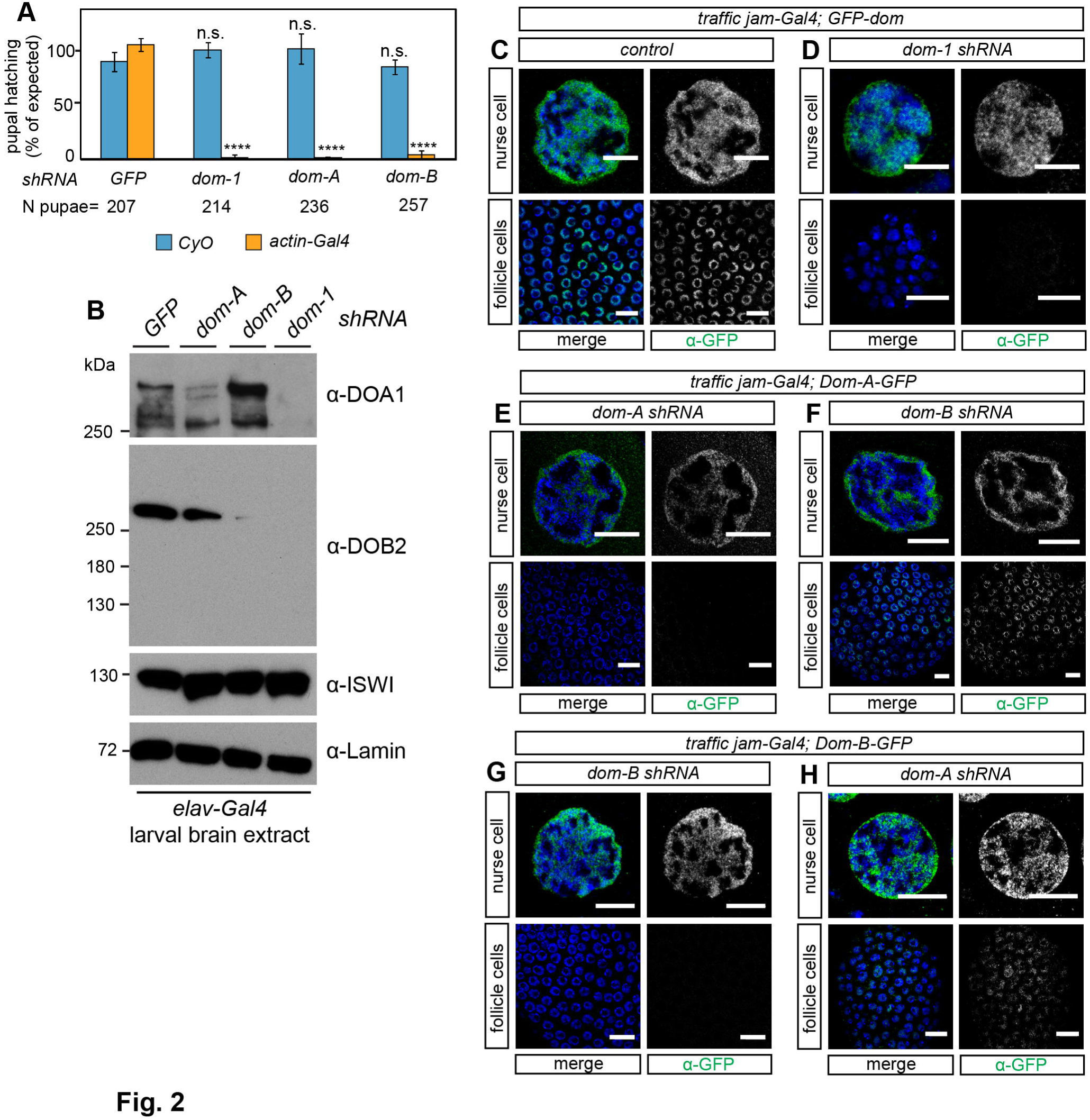
Cell type-specific knockdown of DOM-A and DOM-B reveals requirements during fly development. (A) DOM-A and DOM-B are essential during fly development. *UAS-shRNA* males for *GFP*, *dom-1*, *dom-A* and *dom-B* were crossed with *actin-Gal4*/*CyO* driver females. Pupal hatching was determined as percentage of observed to expected frequency of *CyO* and *actin-Gal4* F1 offspring. The data show mean values of percentage with SD from three biological replicates. N pupae represents the total number of scored pupae. Two-tailed Student’s t-test was used in comparison to *CyO* or *actin-Gal4* siblings with *GFP shRNA*, respectively. **** represents a p-value of <0.0001 and n.s. represents not significant. (B) Validation of specificity for *dom-A* and *dom-B shRNA* knockdowns. *UAS-shRNA* males for *GFP*, *dom-1*, *dom-A* and *dom-B* were crossed with *elav-Gal4* driver females. Larval brains are probed with α-DOA1 17F4 and α-DOB2 4H4 in Western blot. Lamin and ISWI signals are shown as controls. (C-G) Validation of specificity for *dom-A* and *dom-B shRNA* knockdown with transgenic *dom* fosmids. GFP-tagged *dom* transgenes are combined with *dom-1*, *dom-A* and *dom-B shRNA* constructs and somatic follicle cell driver *traffic jam-Gal4*. Representative immunofluorescence images of nurse and follicle cells with staining of GFP (green) and DNA (blue) are shown. Scale bar: 10 μm. Please also refer to Fig. S3A-I.

DOM is well known as a regulator of Notch signaling in wing formation (Eissenberg et al., 2005; Ellis et al., 2015; Gause et al., 2006; Kwon et al., 2013). We also scored typical Notch phenotypes, such as defective wing margins and wing nicks, with high penetrance upon expression of *dom-1* and *dom-2 shRNA* from the *C96-Gal4* driver (Fig. S2). Interestingly, the two DOM isoforms appear to have distinct functions in wing development since *dom-A shRNA* led to wing margin phenotypes with high penetrance, while depleting *dom-B* had only minor effects (Fig. S2D).

The phenotypical RNAi analysis suggested an efficient depletion of DOM-A and DOM-B in different developmental stages. To further document the efficiency and specificity of RNAi depletion we raised monoclonal antibodies against peptides of DOM-A (α-DOA1) and DOM-B (α-DOB2) for Western blot detection. No DOM signals were detected in larval brain extracts with α-DOA1 and α-DOB2 upon *dom-1 shRNA* depletion driven by *elav-Gal4* (Fig. 2B). Furthermore, signals for DOM-A and DOM-B were strongly reduced upon depletion with *dom-A* and *dom-B shRNA*, respectively (Fig. 2B). As a control, levels of the SWI2/SNF2 ATPase Imitation Switch (ISWI) were unaltered (Fig. 2B).

Despite all efforts, the monoclonal DOM-A and DOM-B antibodies could not be used for IFM applications. To circumvent this problem, we combined GFP-tagged *dom* transgenes with *shRNA* constructs expressed from the *traffic jam-Gal4* driver. This driver expresses specifically in somatic follicle cells but not in germline cells such as nurse cells (Olivieri et al., 2010). We detected the *GFP-Dom* transgene in follicle and nurse cells via the GFP antibody (Fig. 2C). We did not detect GFP signal in follicle cell nuclei upon *dom-1*, *dom-A* and *dom-B shRNA* depletion in the respective *GFP-dom*, *dom-A-GFP* and *dom-B-GFP* transgene lines (Fig. 2D,E,G). However, GFP signal was detected in nurse cell nuclei of the same egg chambers (Fig. 2D,E,G), confirming the cell type-specificity of the *traffic-jam* driver. Importantly, DOM-A or DOM-B depletion did not affect the localization of the respective other isoform (Figs. 2F,H and S3), arguing for an independent localization of the two DOM isoforms to chromatin of follicle cells. We conclude that *shRNA* against *dom*, *dom-A* and *dom-B* specifically and efficiently deplete their corresponding target protein.

### Loss of DOM-A in germline cells leads to early cystoblast defects while loss of DOM-B generates defective late egg chambers

Previous analysis could not reveal the specific contributions of DOM-A and DOM-B function to germline development (Ruhf et al., 2001; Yan et al., 2014). We confirmed the importance of DOM for fertility by scoring the egg laying capacity of F1 females from crosses of *shRNA* constructs against *dom*, *dom-A* and *dom-B* with the germline-specific *MTD-Gal4* driver (Fig. 3A) (Yan et al., 2014). We found the egg laying capacity strongly impaired in the absence of DOM, but also upon separate depletion of each isoform (Fig. 3B).

**Fig. 3.**
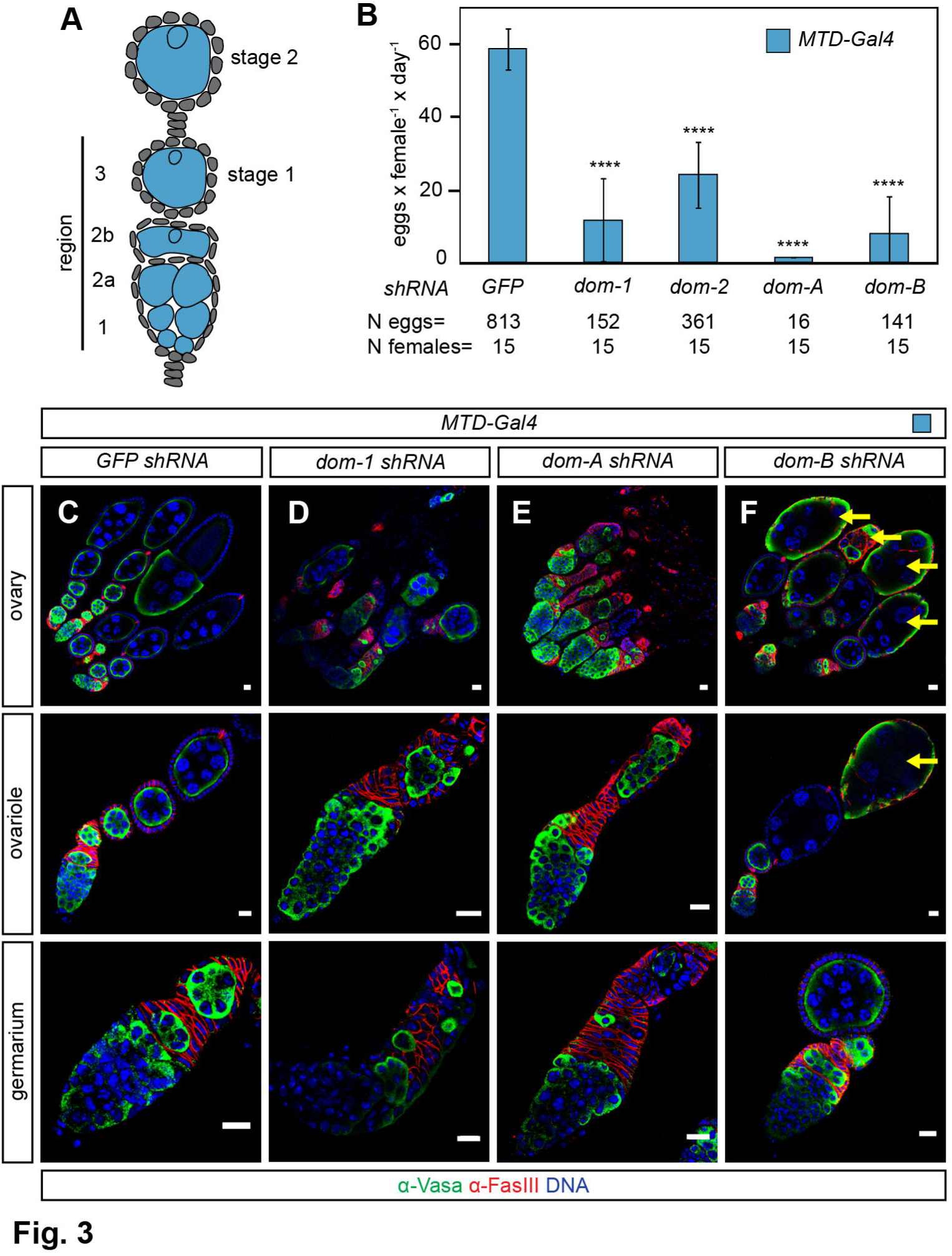
Loss of DOM-A in germline cells leads to early cystoblast defects while loss of DOM-B generates defective late egg chambers. (A) Schematic drawing showing early stages of *D. melanogaster* oogenesis. Specific expression pattern of *MTD-Gal4* driver in germline cells is highlighted in blue. (B) Quantification of egg laying capacity. The data show mean values of laid eggs per female and day with SD of three biological replicates. N eggs represents the total number of scored eggs and N females the total number of analyzed females. Two-tailed Student’s t-test was used in comparison to *GFP shRNA*. **** represents a p-value of <0.0001. (C-F) Immunofluorescence images of ovary, ovariole and germarium with staining of Vasa (green), Fasciclin III (red) and DNA (blue) are shown for the following genotypes: *MTD-Gal4* with (C) *GFP*, (D) *dom-1*, (E) *dom-A* and (F) *dom-B shRNA*. Yellow arrow indicates defective late egg chamber. Scale bar: 10 μm.

We stained ovaries of knockdown animals for germline-specific cytoplasm marker Vasa and follicle cell-specific adhesion molecule Fasciclin III (Fig. 3C-F). As expected, depletion with *dom-1 shRNA* led to an agametic phenotype with characteristic small ovaries and defects in cyst differentiation in the germarium (Fig. 3D) (Yan et al., 2014). Depletion of DOM-A resembled this phenotype (Fig. 3E). In contrast, germline development in the germarium appeared normal upon DOM-B depletion (Fig. 3F), but under these conditions many defective late egg chambers were observed (Fig. 3F). We conclude that the two DOM splice variants are both required for proper oogenesis, but they differ in their requirements: DOM-A is essential for early germline development and DOM-B is critical for egg chamber development at later stages of oogenesis.

### Loss of DOM-A or DOM-B in germline cells from stage 1 onwards leads to ‘dumpless’ egg phenotype

DOM-B was required in germline cells outside of the germarium for egg chamber development. The abundant defects in the germarium upon DOM-A depletion precluded the assessment of a similar role for this isoform during later stages of oogenesis. Therefore, we combined the *mat*α*4-Gal4* driver, which specifically expresses in the germline from stage 1 egg chambers onwards (Fig. 4A) (Yan et al., 2014), with different *dom shRNA* constructs and stained ovaries for Vasa and the cytoplasmic oocyte marker Orb. Notably, *mat*α*4-Gal4*-directed knockdown did not affect the development of germarium and ovariole structures but resulted in defective late egg chambers with short eggs and non-fragmented nurse cells characteristic of the ‘dumpless’ egg phenotype (Fig. 4B-E). Interestingly, individual loss of either DOM-A or DOM-B gave rise to ‘dumpless’ eggs at levels comparable to the combined DOM depletion (Fig. 4F) and accordingly to significantly reduced egg laying (Fig. 4G). Our analysis now shows that both isoforms are required in the germline from stage 1 of oogenesis for the production of proper egg chambers.

**Fig. 4.**
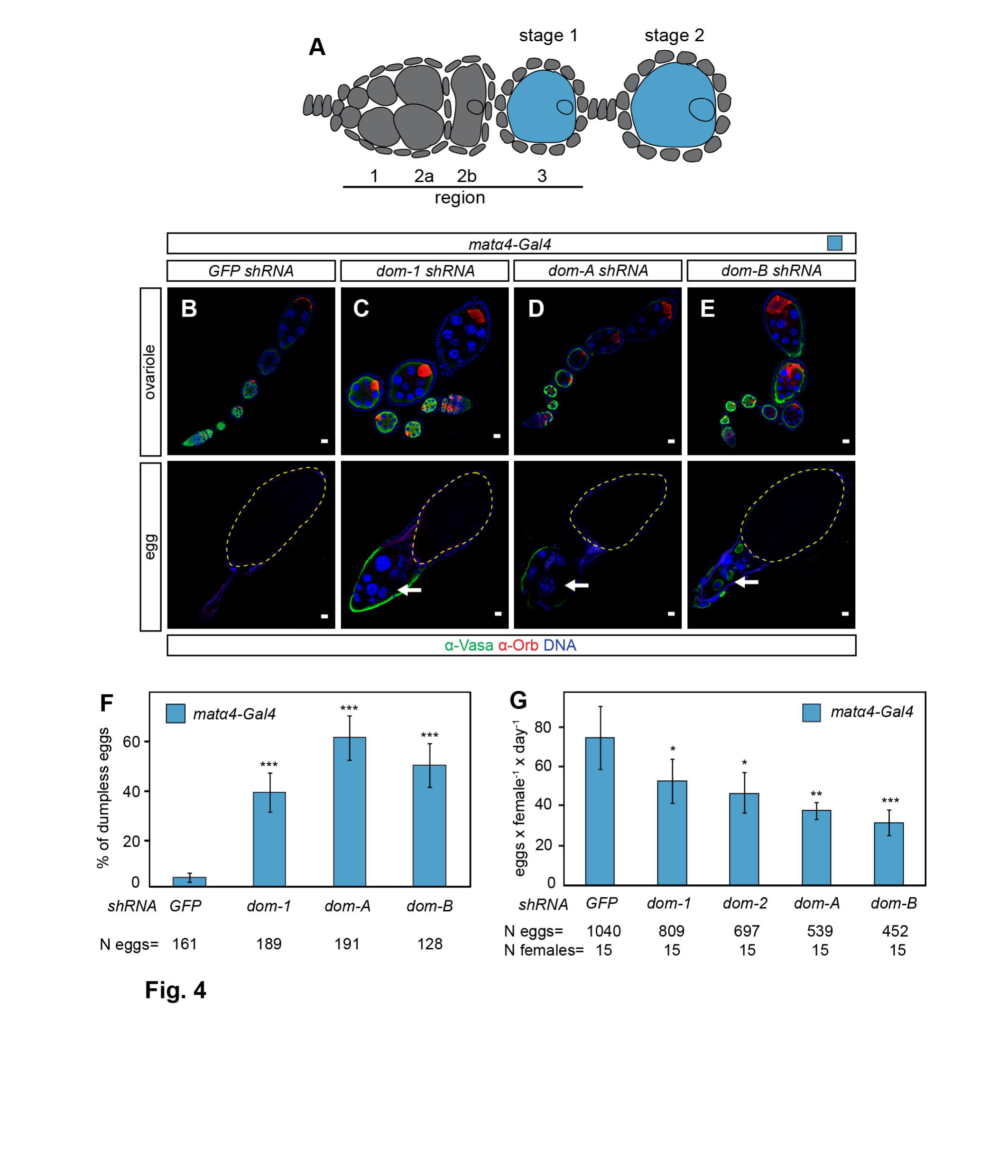
Loss of DOM in germline cells from stage 1 onwards leads to ‘dumpless’ egg phenotype. (A) Schematic drawing showing early stages of *D. melanogaster* oogenesis. Specific expression pattern of *mat*α*4-Gal4* driver in germline cells from stage 1 of oogenesis is highlighted in blue. (B-E) Immunofluorescence images of ovariole and egg with staining of Vasa (green), Orb (red) and DNA (blue) are shown for the following genotypes: *mat*α*4-Gal4* with (B) *GFP*, (C) *dom-1*, (D) *dom-A* and (E) *dom-B shRNA*. White arrow indicates anterior end of egg chamber with non-fragmented nurse cells. Yellow dashed line indicates posterior egg. Scale bar: 10 μm. (F) Quantification of ‘dumpless’ egg phenotype. The data show mean values in percentage with SD of three biological replicates. N eggs represents the total number of scored eggs. Two-tailed Student’s t-test was used in comparison to *GFP shRNA*. *** represents a p-value of <0.001. (G) Quantification of egg laying capacity. The data show mean values of laid eggs per female and day with SD of three biological replicates. N eggs represents the total number of scored eggs and N females the total number of analyzed females. Two-tailed Student’s t-test was used in comparison to *GFP shRNA*. * represents a p-value of <0.05, ** <0.01 and *** <0.001.

### Loss of DOM-A or DOM-B in somatic follicle cells leads to severe packaging defects

Domino function is not only required in the germline but is also essential for somatic follicle cell development (Xi and Xie, 2005). We wondered whether both DOM splice variants were required for follicle cell function. To address this, we used again the *traffic jam-Gal4* driver (Fig. 5A), in combination with different *dom shRNA* constructs. We scored severe packaging phenotypes upon DOM, DOM-A or DOM-B depletion ranging from two cysts in a single egg chamber to complete ovariole fusions (Fig. 5B-E). Furthermore, many germaria showed additional α-Spectrin positive stalk-like structures without germline cysts (Fig. 5F-I) indicating defects in the coordination of follicle cell proliferation with cyst differentiation. Remarkably, we observed egg chambers with abnormal cyst numbers from one cell to more than sixteen cells (Fig. 5B-E) suggesting a soma-dependent proliferation defect in the germline. Defective late egg chambers were found in the process of decay as verified by staining for activated caspase-3 (Fig. 5J-M). As a result, egg laying capacity was drastically impaired with two different *dom shRNA*s as well as with *dom-A* and *dom-B shRNA* (Fig. 5N).

**Fig. 5.**
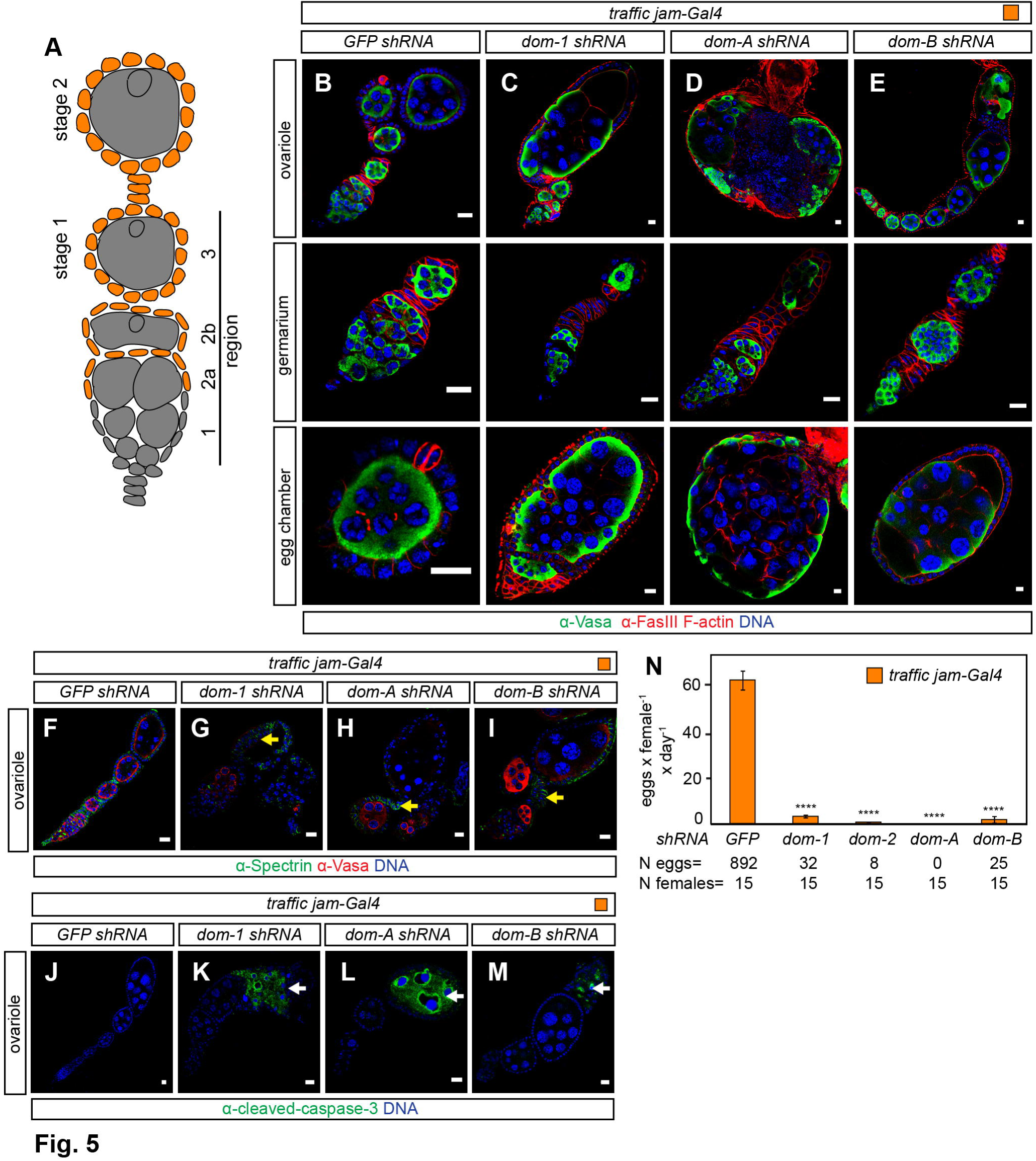
Loss of DOM-A or DOM-B in somatic follicle cells leads to severe packaging defects. (A) Schematic drawing showing early stages of *D. melanogaster* oogenesis. Specific expression pattern of *traffic jam-Gal4* driver in somatic follicle cells is highlighted in orange. (B-E) Immunofluores-cence images of ovariole, germarium and egg chamber with staining of Vasa (green), Fasciclin III and F-actin (red) and DNA (blue) are shown for the following genotypes: *traffic jam-Gal4* with (B) *GFP*, (C) *dom-1*, (D) *dom-A* and (E) *dom-B shRNA*. Scale bar: 10 μm. (F-I) Immunofluorescence images of ovarioles with staining of Spectrin (green), Vasa (red) and DNA (blue) are shown for the following genotypes: *traffic jam-Gal4* with (A) *GFP*, (B) *dom-1*, (C) *dom-A* and (D) *dom-B shRNA*. Yellow arrow indicates stalk-like structures. Scale bar: 10 μm. (J-M) Immunofluorescence images of ovarioles with staining of Cleaved Caspase-3 (green) and DNA (blue) are shown for the following genotypes: *traffic jam-Gal4* with (A) *GFP*, (B) *dom-1*, (C) *dom-A* and (D) *dom-B shRNA*. White arrow indicates apoptotic egg chamber. Scale bar: 10 μm. (N) Quantification of egg laying capacity. The data show mean values of laid eggs per female and day with SD of three biological replicates. N eggs represents the total number of scored eggs and N females the total number of analyzed females. Two-tailed Student’s t-test was used in comparison to *GFP shRNA*. **** represents a p-value of <0.0001.

We validated packaging phenotypes and sterility with another somatic driver line, *c587-Gal4*, which expresses in somatic escort cells and early follicle cells in the germarium (Fig. S4A) (Eliazer et al., 2011; Kai and Spradling, 2003). Staining with the oocyte marker Orb revealed the nature of packaging defects as compound egg chambers with oocytes at opposite positions of an egg chamber and confirmed apoptotic phenotype in late egg chambers (Fig. S4B-E). We conclude that DOM-A and DOM-B function is critical in somatic follicle cells for proper packaging of egg chambers.

### DOM-B promotes global H2A.V incorporation into germline chromatin of the germarium

We wished to explore roles of DOM isoforms for H2A.V incorporation into chromatin and γH2A.V turnover *in vivo*. Towards this end, we raised a specific polyclonal antibody against a H2A.V peptide. Staining ovarioles with this antibody and a previously characterized γH2A.V antibody (Lake et al., 2013) resembled the published H2A.V localization pattern (Fig. 6A-D) (Joyce et al., 2011). We found H2A.V ubiquitously expressed in somatic and germline cells of the germarium with a notable enrichment on GSCs (Fig. 6B,L). Remarkably, H2A.V and γH2A.V signals were only detected in nurse cells up to stage 5 of oogenesis (Fig. 6C), while both signals were absent in later nurse cell nuclei (Fig. 6D). However, H2A.V was present in the surrounding follicle cells at all stages, providing convenient staining controls (Fig. 6A-D). These observations are in general agreement with previous studies (Joyce et al., 2011), except that we did not detect signals for H2A.V in oocytes, possibly due to differences in H2A.V epitope sequence.

**Fig. 6.**
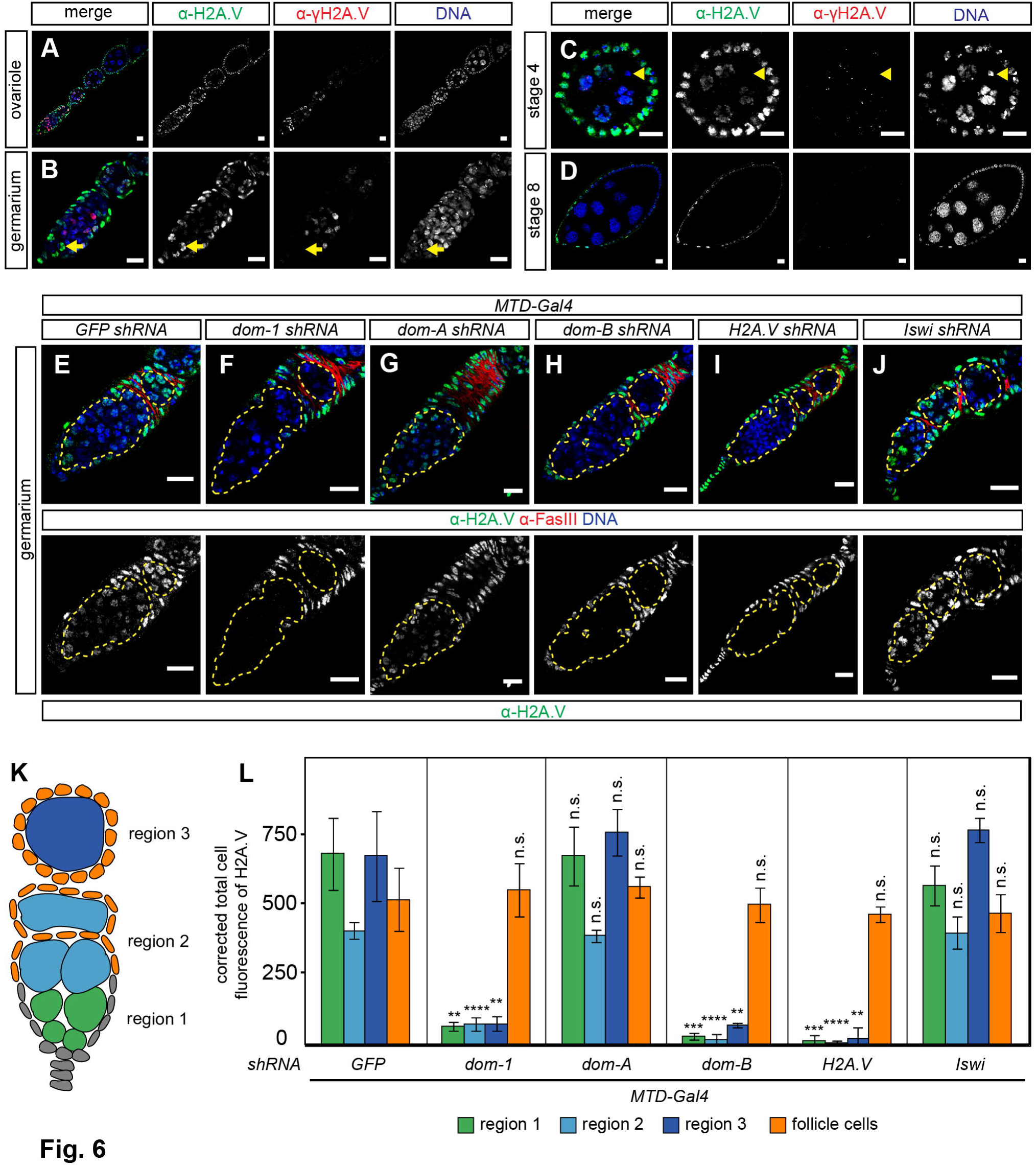
DOM-B promotes global H2A.V incorporation into germline chromatin of the germarium. (A-D) Immunofluorescence images of ovariole, germarium, stage 4 and 8 egg chambers with staining of H2A.V (green), γH2A.V (red) and DNA (blue) are shown for wild type. Yellow arrow and arrowhead indicate GSCs and oocyte, respectively. Scale bar: 10 μm. (E-J) Immunofluorescence images of ovariole with staining of H2A.V (green), Fasciclin III (red) and DNA (blue) are shown for the following genotypes: *MTD-Gal4* with (E) *GFP*, (F) *dom-1*, (G) *dom-A*, (H) *dom-B*, (I) *H2A.V* and (J) *Iswi shRNA*. Yellow dashed line indicates germline cells. Scale bar: 10 μm. (K) Schematic drawing of *D. melanogaster* germarium. Green, light and dark blue indicate region 1, 2 and 3 of the germarium, respectively. (L) Quantification of H2A.V signals in germline and somatic cells of the germarium. Corrected total cell fluorescence (CTCF) of H2A.V was calculated for 30 germline cysts in different regions of the germarium and follicle cells, respectively. Mean CTCF values with SD of three biological replicates are shown. Color code represents cell types in Fig. 6K. Two-tailed Student’s t-test was used in comparison to *GFP shRNA*. ** represents a p-value of <0.01, *** <0.001 and **** <0.0001 and n.s not significant.

Upon depletion of DOM and DOM-B with *MTD-Gal4* we found that H2A.V was lost specifically in germline cells of the germarium (Fig. 6F,H,K,L). H2A.V was essentially absent since the signals were comparable to *H2A.V shRNA* (Fig. 6I,K,L). As a control, levels of H2A.V were unaltered in somatic follicle cells, documenting the germline-specificity of the effect (Fig. 6E-L). In remarkable contrast, loss of DOM-A did not affect levels of H2A.V (Fig. 6G,K and L). Likewise, depletion of ISWI ATPase, which causes similar germline phenotypes (Ables and Drummond-Barbosa, 2010; Xi and Xie, 2005; Yan et al., 2014), also did not affect H2A.V levels (Fig. 6J,K and L). We conclude that DOM-B, but not DOM-A, is required for global incorporation of H2A.V into germline chromatin of the germarium.

### DOM-independent H2A.V incorporation in germline nurse cells

To monitor H2A.V levels in germline cells outside of the germarium we again employed the *mat*α*4-Gal4* driver, which specifically expresses from stage 1 of oogenesis onwards (Yan et al., 2014). As expected, *mat*α*4-Gal4-*directed knockdown of DOM, DOM-A and DOM-B did not affect H2A.V levels in the germarium (Figs. S5,S6). As a control, *H2A.V shRNA* knockdown specifically depleted H2A.V and γH2A.V in germline nurse cells of stage 3 egg chambers (Fig. 7A,E and F). Surprisingly however, H2A.V and γH2A.V was unaltered in germline nurse cells up to stage 4 egg chambers upon DOM, DOM-A or DOM-B knockdown (Figs 7A-D and F, S5 and S6). Notably, depletion of another SWI/SNF remodeler, Inositol-requiring protein 80 (dINO80), with two different *dIno80 shRNAs* and drivers did not affect H2A.V localization in ovarioles (Fig. S7). Although, a previous study suggests efficient knockdown since RNAi resembles embryonic lethality phenotype of *dIno80* mutants (Bhatia et al., 2010; Yan et al., 2014). This observation suggests that H2A.V incorporation into germline chromatin from stage 2 onwards might be independent of Domino.

**Fig. 7.**
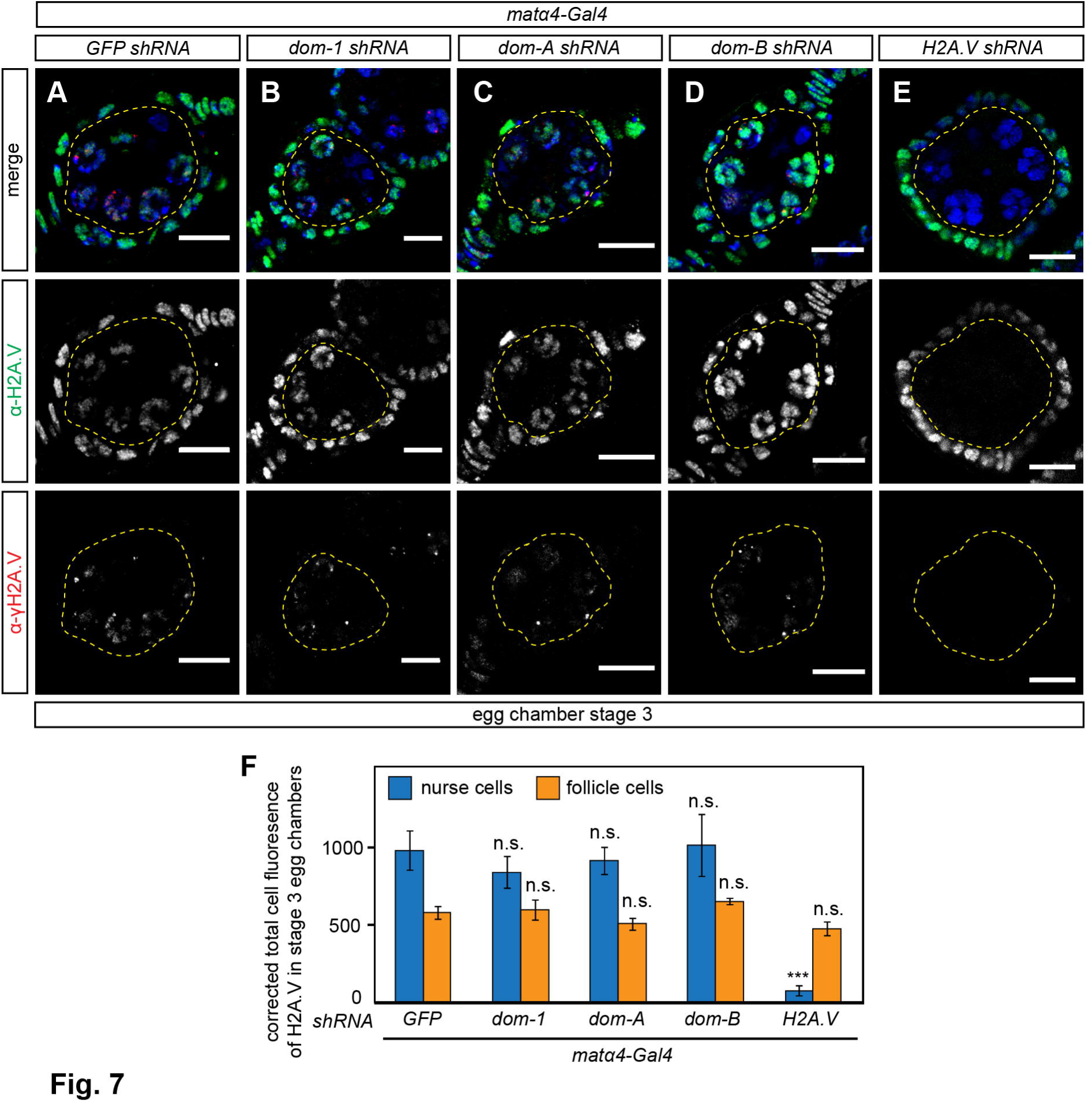
DOM-independent H2A.V incorporation in germline nurse cells. (A-E) Immunofluorescence images of egg chambers with staining of H2A.V (green), γH2A.V (red) and DNA (blue) are shown for the following genotypes: *mat*α*4-Gal4* with (A) *GFP*, (B) *dom-1*, (C) *dom-A*, (D) *dom-B* and (E) *H2A.V shRNA*. Yellow dashed line indicates germline nurse cells. Scale bar: 10 μm. (F) Quantification of H2A.V signals in nurse and follicle cells of stage 3 egg chambers. Corrected total cell fluorescence (CTCF) of H2A.V was calculated for 50 germline nurse cells and somatic follicle cells, respectively. Mean CTCF values with SD of three biological replicates are shown. Two-tailed Student’s t-test was used in comparison to *GFP shRNA*. *** represents a p-value of <0.001 and n.s. not significant.

### Involvement of DOM-A in H2A.V eviction in germline cells from stage 5 onwards

We next assessed how depletion of individual DOM isoforms affect global H2A.V and γH2A.V in germline nurse cells of later egg chamber stages. Remarkably, we found that H2A.V and γH2A.V signals persisted in stage 8 egg chambers upon DOM or DOM-A depletion (Figs. 8B,C,G,H,K, S5 and S6). By contrast, H2A.V was removed in the absence of DOM-B as in wild type ovarioles (Figs 8A,E,F,I,K, S5 and S6). Evidently, the DOM-A isoform is specifically involved in removing H2A.V from germline cells by stage 5 of oogenesis, illustrating once more the functional diversification of the two splice variants.

**Fig. 8.**
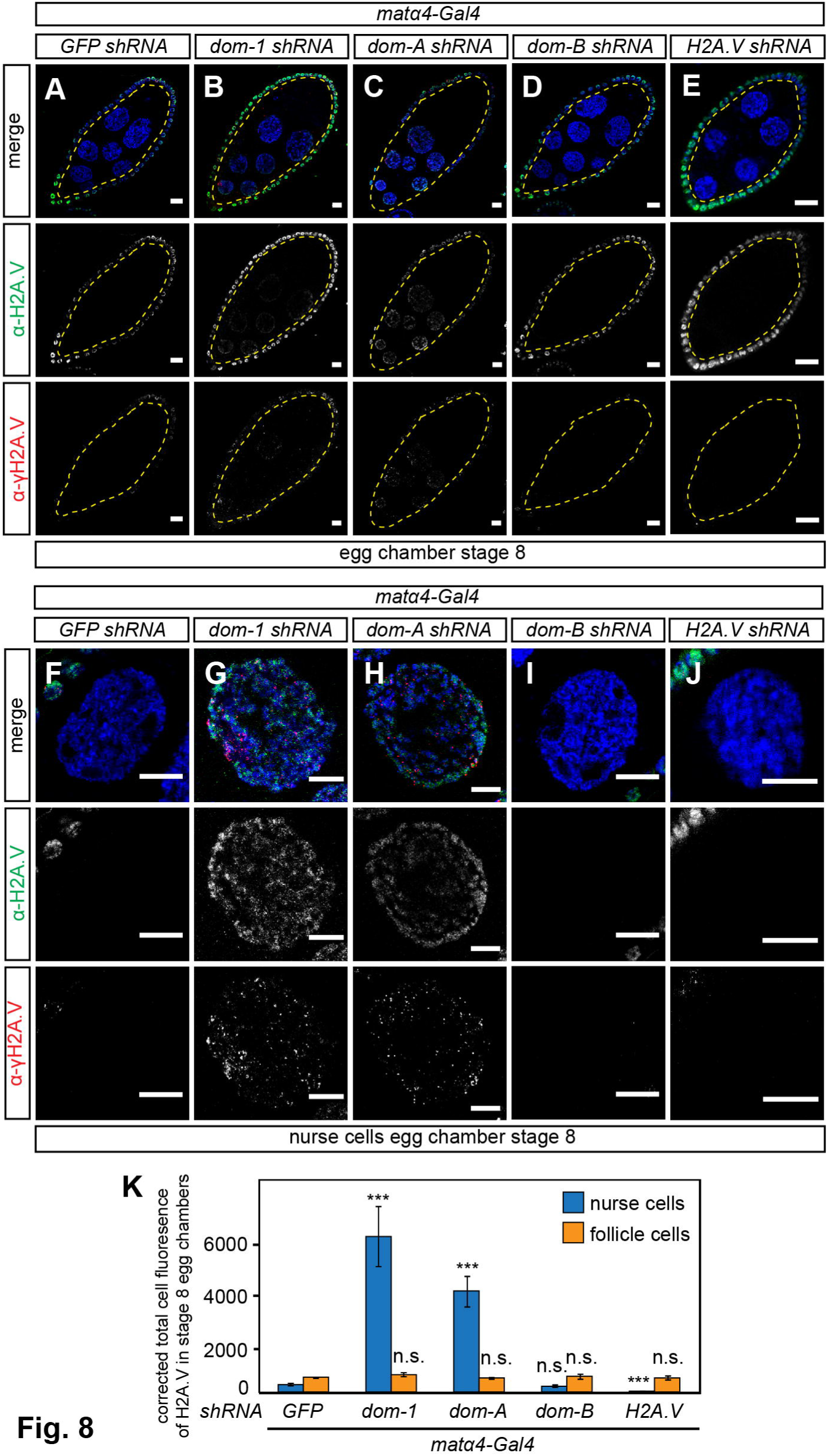
Involvement of DOM-A in H2A.V eviction in germline cells from stage 5 onwards. (A-J) Immunofluorescence images of stage 8 egg chambers and nurse cells with staining of H2A.V (green), γH2A.V (red) and DNA (blue) are shown for the following genotypes: *mat*α*4-Gal4* with (A, F) *GFP*, (B, G) *dom-1*, (C, H) *dom-A*, (D, I) *dom-B* and (E, J) *H2A.V shRNA*. Yellow dashed line indicates germline nurse cells. Scale bar: 10 μm. (K) Quantification of H2A.V persistence in germline nurse cells of stage 8 egg chambers. Corrected total cell fluorescence (CTCF) of H2A.V was calculated for 50 germline nurse cells and somatic follicle cells, respectively. Mean CTCF values with SD of three biological replicates are shown. Two-tailed Student’s t-test was used in comparison to *GFP shRNA*. *** represents a p-value of <0.001 and n.s. not significant.

### DOM-B promotes global H2A.V incorporation into dividing somatic follicle cell chroma-tin

To address whether other cell types also utilize DOM for H2A.V incorporation, we used the somatic follicle cell driver *traffic jam-Gal4* for knockdown in all stages of follicle cell development. Like in germline cells of the germarium, we observed that depletion of DOM and DOM-B led to complete absence of H2A.V signals in follicle cell nuclei (Fig. A,B,D,G,H), when compared to the corresponding *H2A.V* knockdown (Fig. 9E,G,H). As before, *dom-A shRNA* did not affect H2A.V levels in follicle cells (Fig. 9C,G,H) and neither did depletion of ISWI (Fig. 9F,G,H), which causes similar packaging phenotypes (Börner et al., 2016). Furthermore, canonical histone H2A (Fig. 9I,J) and heterochromatin-associated HP1 protein (Fig. 9K,L) were unaltered upon DOM depletion indicating a globally intact chromatin structure in the absence of H2A.V.

**Fig. 9.**
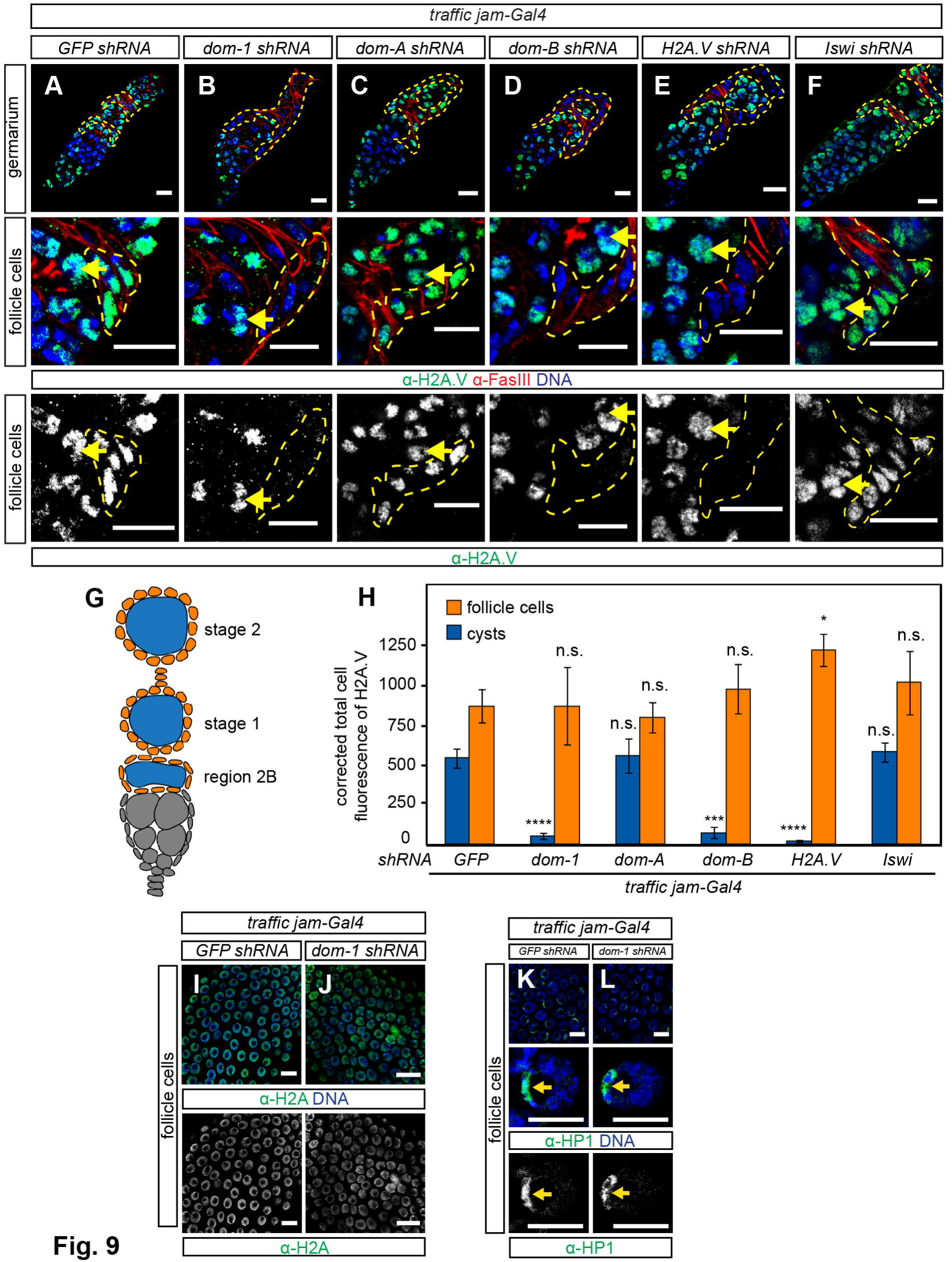
M-B promotes global H2A.V incorporation into somatic follicle cell chromatin. (A-F) Immunofluorescence images of germarium with staining of H2A.V (green), Fasciclin III and F-actin (red) and DNA (blue) are shown for the following genotypes: *traffic jam-Gal4* with (A) *GFP*, (B) *dom-1*, (C) *dom-A*, (D) *dom-B*, (E) *H2A.V* and (F) *Iswi shRNA*. Yellow dashed line indicates somatic follicle cells. Yellow arrow indicate germline cysts. Scale bar: 10 μm. (G) Schematic drawing showing early stages of *D. melanogaster* oogenesis. Orange and blue indicate follicle cells and cysts, respectively, that were used for measuring H2A.V signal intensities in Fig. 8H. (H) Quantification of H2A.V signals in somatic and germline cells. Corrected total cell fluorescence (CTCF) of H2A.V was calculated for 30 somatic follicle cells and cysts/nurse cells, respectively. Mean CTCF values with SD of three biological replicates are shown. Color code represents cell types in Fig. 8G. Two-tailed Student’s t-test was used in comparison to *GFP shRNA*. * represents a p-value of <0.05, *** <0.001 and **** <0.0001 and n.s not significant. (I, J) Immunofluorescence images of follicle cells with staining of H2A (green) and DNA (blue) are shown for the following genotypes: *traffic jam-Gal4* with (I) *GFP* and (J) *dom-1 shRNA*. Scale bar: 10 μm. (K, L) Immunofluorescence images of follicle cells with staining of HP1 (green) and DNA (blue) are shown for the following genotypes: *traffic jam-Gal4* with (K) *GFP* and (L) *dom-1 shRNA*. Scale bar: 10 μm.

At the anterior end of the germarium a subset of non-dividing somatic cells, the terminal filament and cap cells, form the stem cell niche. These terminal differentiated cells highly expressed H2A.V (Fig. S8B). The *hedgehog-Gal4* (*hh*) driver specifically expresses in somatic filament and cap cells (Fig. S8A) (Eliazer et al., 2011; Forbes et al., 1996; Pan et al., 2007), permitting an assessment of DOM function in these cells. Surprisingly, H2A.V signals in niche cells were not changed upon DOM, DOM-A or DOM-B depletion (Fig. S8B-E), although *hh*-mediated knockdown of DOM or DOM-A caused pupal lethality at 29°C (data not shown). These findings document a cell type-specific requirement for the DOM-B isoform for H2A.V chromatin incorporation in dividing follicle cells but not in terminal differentiated niche cells.

In summary, our systematic analysis showed that nucleosome remodeling factor isoforms DOM-A and DOM-B have non-redundant functions in germline and soma in the formation of egg chambers. We further demonstrated that the two DOM isoforms have distinct functions in H2A.V exchange during *D. melanogaster* oogenesis.

## Discussion

In *D. melanogaster* the properties of the two ancient, ubiquitous histone H2A variants, H2A.X and H2A.Z, are combined in a single molecule, H2A.V (Baldi and Becker, 2013; Talbert and Henikoff, 2010). Given that H2A.V carries out the very different functions of the two former variants as DNA damage sensor and architectural element of active promoters (Madigan et al., 2002; Mavrich et al., 2008), and its further roles in heterochromatin formation (Chioda et al., 2010; Hanai et al., 2008; Qi et al., 2006), this histone appears loaded with regulatory potential. Accordingly, placement of the variant, either randomly along with canonical H2A at replication, or more specifically by exchange of canonical H2A through nucleosome remodeling factors becomes a crucial determinant for function.

Replacement of H2A-H2B dimers with variants is catalyzed by dedicated nucleosome remodeling factors. Mechanistic detail comes from the analysis of the yeast SWR1 complex, which incorporates H2A.Z at strategic positions next to promoters (Luk et al., 2010; Mizuguchi, 2004; Ranjan et al., 2013). Sequence similarity searches revealed only one SWR1 homolog in fruit flies, the Domino (DOM) protein (Ruhf et al., 2001). Phenotypes of mutant *domino* alleles suggest broad requirements for cell proliferation and differentiation, homeotic gene silencing, Notch signaling, hematopoiesis and oogenesis (Braun et al., 1998; Eissenberg et al., 2005; Ellis et al., 2015; Kwon et al., 2013; Lu et al., 2007; Ruhf et al., 2001; Sadasivam and Huang, 2016; Walker et al., 2011; Xi and Xie, 2005; Yan et al., 2014). These requirements are presumed to be due to a wide-spread involvement of Domino in transcription regulation. H2A.V-containing nucleosomes demarcate a series of well-positioned nucleosomes just downstream of most transcription start sites and are important for transcription initiation (Mavrich et al., 2008). Our comprehensive loss-of-function analysis using the oogenesis example reveals that most H2A.V incorporation depends on Domino.

So far the published phenotypes associated with *dom* mutant alleles have not been systematically complemented (Braun et al., 1997; Eissenberg et al., 2005; Ruhf et al., 2001). Our comprehensive complementation analysis shows that *dom* mutant phenotypes are indeed due to defects in the *dom* gene. Remarkably, *dom* lethality and sterility can be partially rescued by complementation with the orthologous human SRCAP gene, providing an impressive example of functional conservation of SWR1-like remodelers (Eissenberg et al., 2005). The contributions of the two splice variants, DOM-A and DOM-B, had not been previously assessed. We now demonstrate that both isoforms are essential for fly development suggesting non-redundant functions that map to the different C-termini. The long DOM-A isoform contains a SANT domain followed by sequences that stand out due to their abundance of glutamines. Q-rich stretches are widely found in modulators of transcriptional regulation (Gemayel et al., 2015). These features are also present in the C-terminus of p400, the second human SWR1 ortholog, but do not occur in either DOM-B or SCRAP (Eissenberg et al., 2005). These findings lead us to speculate that distinct functions of p400 and SCRAP in humans may be accommodated to some extent by the two splice forms of Domino in flies. Accordingly, it will be interesting to explore whether the two isoforms reside in distinct complexes. This may well be the case since previous affinity purification of a Tip60 complex subunit using a tagged pontin apparently only identified DOM-A, but not DOM-B (Kusch et al., 2004). Remarkably, the SANT domain of p400 interacts directly with TIP60 and inhibits its catalytic activity (Park et al., 2010) providing an interesting lead for further investigation of DOM splice variants and TIP60 interactions.

Following up on the initial observation of early defects in germline stem cells and cyst differentiation upon loss of both DOM isoforms (Yan et al., 2014), we now find that this phenotype is exclusively caused by loss of the DOM-A isoform. Interestingly, studies with human embryonic stem cells show that p400/TIP60 integrates pluripotency signals to regulate gene expression (Chen et al., 2013; Fazzio et al., 2008) suggesting similar roles for DOM-A in germline stem cells. This is in contrast with requirements for both isoforms for germline development in later stages of oogenesis, highlighting a developmental specialization of the two DOM remodelers.

DOM is also involved in the differentiation and function of somatic cells in the germarium. It had been known that *dom* mutant SSC clones show defects in self-renewal leading to their subsequent loss (Xi and Xie, 2005). Our data now document non-redundant requirements of both DOM isoforms in somatic cells for proper coordination of follicle cell proliferation with cyst differentiation. Failure to adjust these two processes leads to 16-cell cyst packaging defects that manifest as compound egg chamber phenotypes. These rare phenotypes had only been described upon perturbation of some signaling pathways such as Notch signaling or Polycomb regulation (Jackson and Blochlinger, 1997; Narbonne et al., 2004). Our findings show essential functions of both DOM isoforms in soma development affecting fly fertility.

We assume that many of the cell specification defects we scored upon DOM depletion are due to compromised H2A.V incorporation, depriving critical promoters of the H2A.Z-related architectural function. Alternatively, scaffolding activities might partially explain some roles of chromatin remodelers as suggested for the transcriptional activation potential of SRCAP (Bowman et al., 2011). Although we cannot rule out former scenario, we favor the idea that observed phenotypes are linked to compromised H2A.V exchange. So far, our knowledge of the mechanisms of H2A.V incorporation has been anecdotal. Roles for Domino in H2A.V incorporation close to the *E2f1* or *hsp70* gene promoters have been described (Kusch et al., 2014; Lu et al., 2007). In mutant germline clones for MRG15, a TIP60/DOM-A complex subunit, H2A.V and γH2A.V was lost in germaria (Joyce et al., 2011). Our comprehensive analysis in the germline and soma revealed a specific involvement for the short DOM-B isoform for the chromatin incorporation of H2A.V at the global level. Given the requirement for both isoforms for cell specification during oogenesis, we speculate that the DOM-B isoform may serve to incorporate bulk H2A.V into chromatin, while the long DOM-A isoform would be more involved in regulatory refinement of location.

While the failures in cell specification and egg morphogenesis are likely to be explained by loss of the H2A.Z-related features of H2A.V, ablation of DOM may also compromise the DNA damage response, which involves phosphorylation of H2A.V (γH2A.V). Conceivably, the role of γH2A.V as a DNA damage sensor may be best fulfilled by a broad distribution of H2A.V throughout chromatin (Baldi and Becker, 2013). Such an untargeted incorporation of the variant may be best achieved by stochastic, chaperone-mediated incorporation during replication (Li et al., 2012), or by an untargeted activity of DOM-B. We observed an example for a DOM-independent incorporation in polyploid nurse cells of stage 2 to 5 egg chambers, in which abundant H2A.V and γH2A.V signals were not affected by depletion of DOM. Polyploidy arises as cells switch their cell cycle program to endoreplication. DOM-independent incorporation of H2A.V may serve to cope with many naturally occurring DNA double-strand breaks during massive endoreplication of nurse cells.

There is isolated evidence that nucleosome remodelers do not only incorporate H2A variants, but can also remove them. In yeast, the genome-wide distribution of H2A.Z appears to be established by the antagonistic functions of SWR1 and Ino80 remodeling complexes, where Ino80 replaces stray H2A.Z-H2B with canonical H2A-H2B dimers (Papamichos-Chronakis et al., 2011). In *D. melanogaster*, a TIP60/DOM-A complex was shown to be involved in the exchange of γ-H2A.V by unmodified H2A.V during the erasure of the DNA damage signal (Kusch et al., 2004). Furthermore, it has been speculated that H2A.V and γH2A.V could be actively removed from nurse cells in late egg chambers, since corresponding signals are absent from stage 5 onwards (Jang et al., 2003; Joyce et al., 2011). We now demonstrate that depletion of DOM-A leads to persistence of H2A.V and γ-H2A.V in nurse cells of late egg chambers, clearly documenting the ability of the remodeler to remove bulk H2A.V and variant modified during DNA damage induction. Our findings highlight the specific requirements of DOM splice variants for incorporation and removal of H2A.V during *D. melanogaster* oogenesis. It remains an interesting and challenging question how DOM-A and DOM-B complexes are targeted genome-wide and function *in vivo* to establish specific H2A.V patterns in different cell types during development.

## Materials and methods

### *D. melanogaster* strains and genetics

The *domino* fosmid derivatives are based on the fosmid library clone pflyfos016675 (kind gift of P. Tomancak, MPI-CPG, Dresden, Germany). The genomic region of *domino* was modified by recombineering in *Escherichia coli* using the pRedFLP4 recombination technology (Ejsmont et al., 2009). All oligonucleotides and their combinations are listed in Supplementary information (Tables S1 and S2). The *GFP-dom* fosmid codes for *domino* with an N-terminal 2xTY1-sGFP-3xFLAG-tag which tags both isoforms *dom-A* and *dom-B*. The *dom-A-GFP* and *dom-B-GFP* fosmids code for *domino* with a C-terminal 2xTY1-sGFP-3xFLAG-tag which tags *dom-A* and *dom-B*, respectively. The entire *domino* locus was replaced with 2xTY1-EGFP-3xFLAG-tag in the Δ*dom-GFP* fosmid. All *domino* fosmid variants were verified by restriction enzyme digestion, PCR and sequencing before injection into *D. melanogaster*. A detailed description of the protocol can be obtained from the authors. Transgenic flies were made by phiC31 integrase-mediated site-specific integration into attP2 landing site in the 3^rd^ chromosome (Genetic Services, Inc., USA). Fosmid constructs contain a *dsRed* cassette driven by 3xP3 promoter to select for transformants. All fosmid, *UAS-shRNA* and *Gal4* driver lines are listed in Supplementary information. For ovary analysis, *UAS-shRNA* males were crossed with *Gal4* driver virgins at 29°C and 5-7 day old F1 females were used, unless not stated otherwise.

### Complementation of lethality phenotype in *dom*^*1*^ and *dom*^*9*^ alleles

Males and female virgins with *dom*^*1*^/*bcg*; *domino*-/+ or *dom*^*9*^/*bcg*; *domino* fosmid/+ were crossed with each other, respectively. F1 offspring were analyzed for *bcg* phenotype and rescue of pupal lethality was determined as percentage of observed to expected frequency of homozygous *dom*^*1*^ or *dom*^*9*^ offspring. Mean values in percentage with SD from three biological replicates were calculated.

### Complementation of necrotic lymph glands in *dom*^*1*^ allele

Necrotic lymph glands were scored in 3^rd^ instar larvae of homozygous *w*^*1118*^, *dom*^*1*^, *dom*^*1*^;*GFP-dom*, *dom*^*1*^;*dom-A-GFP* and *dom*^*1*^;*dom-B-GFP*. Mean values in percentage with SD from three biological replicates were calculated.

### Complementation of sterility phenotype in *dom*^*9*^ allele

Homozygous female virgins for *w*^*1118*^, *dom*^*9*^, *dom*^*9*^;*GFP-dom*, *dom*^*9*^;*dom-A*, *dom*^*9*^;*dom-B* and *dom*^*9*^;Δ*dom-GFP* were collected for 2-3 days at 25°C. Females of each genotype were mated with *w*^*1118*^ males in vials with yeast paste for 2 days. Females were put in individual vials for 3 days without males and laid eggs were counted daily. The egg laying capacity was determined as the number of laid eggs per female and day. The data was averaged and mean values with SD from three biological replicates were calculated.

### Pupal hatching assay

*UAS-shRNA* males for *GFP*, *dom-1*, *dom-A* and *dom-B* were crossed with *actin-Gal4*/*CyO* driver virgins at 25°C. Pupae were transferred to new vials and numbered. A 1:1 ratio of *CyO* and *actin-Gal4* F1 siblings was expected. *CyO* phenotype was counted and pupal hatching was determined as percentage of observed to expected frequency of *CyO* and *actin-Gal4* F1 siblings. Mean values in percentage with SD from three biological replicates were calculated.

### Analysis of wing phenotypes

*UAS-shRNA* males for *GFP*, *dom-1*, *dom-2, dom-A* and *dom-B* were crossed with *C96-Gal4* driver virgins at 29°C. Wing phenotypes were scored in F1 offspring and relative mean values in percentages with SD from three biological replicates were calculated.

### Egg laying assay

Homozygous *UAS-shRNA* males for *GFP*, *dom-1*, *dom-2, dom-A* and *dom-B* were crossed with *MTD*, *mat*α*4*, *c587* or *traffic jam-Gal4* driver virgins at 29°C. F1 female virgins of each genotype (2-3 days old) were mated with *w*^*1118*^ males in vials with yeast paste for 2 days. Females were then put in individual vials without males and laid eggs were counted daily. The egg laying capacity was determined as the number of laid eggs per female and day. The data was averaged and mean values with SD from three biological replicates were calculated.

### Generation of DOA1, DOB2 and H2A.V antibodies

DOA1 (KEHKRSRTDAGYDGSRRPNC) and DOB2 (TPKESQSEPRRKITQPKC) peptides with N-terminal PEG-Biotin and C-terminal coupled Ovalbumin were made by Peptide Specialty Laboratories (PSL, Heidelberg, Germany). The monoclonal rat α-DOA1 17F4 and mouse α-DOB2 4H4 peptide antibodies were developed by E. Kremmer (Helmholtz Zentrum Munich, Germany). The specificity was confirmed in Western blot by RNAi (Fig. 2B) and with recombinantly expressed DOM-A and DOM-B (data not shown).

H2A.V peptide (QPDQRKGNVIL) corresponding to aa 127-137 of H2A.V was coupled to KLH via C-terminal cysteine and used to raise polyclonal rabbit α-H2A.V antibody (Eurogentec, Netherlands). The specificity of α-H2A.V was validated by loss of H2A.V immunofluorescence signal upon cell type-specific knockdown with *H2A.V shRNA* (Figs 6I,L, 7E,F, 8E,H).

### Immunological techniques and microscopy

Immunofluorescence microscopy was performed using standard procedures and the Leica TCS SP5 II confocal microscope. All antibodies are listed in Supplementary information. Images were processed using ImageJ (NIH, USA) and Adobe Photoshop. Please refer to Supplementary information for more details.

### Western blot

Twelve pairs of ovaries or 10 brains of 3^rd^ instar larvae were dissected, homogenized in Laemmli buffer with a pestle and incubated at 95°C for 5 minutes. Western blot was performed using standard procedures and all antibodies are listed in Supplementary Information.

## Acknowledgements

Stocks obtained from the Bloomington Drosophila Stock Center (NIH P40OD018537) were used in this study. We thank the TRiP at Harvard Medical School (NIH/NIGMS R01-GM084947) for providing transgenic RNAi fly lines. The monoclonal antibodies Orb 4H8 and 6H4 were developed by Paul Schedl, 7G10 anti-Fasciclin III was developed by Corey Goodman, 3A9 (323 or M10-2) was developed by Branton, D. and Dubreuil, R., C1A9 was developed by Wallrath, L.L. and UNC93-5.2.1 was developed by R. Scott Hawley. They were obtained from the Developmental Studies Hybridoma Bank, created by the NICHD of the NIH and maintained at The University of Iowa, Department of Biology, Iowa City, IA 52242.

We thank A. Eberharter, J-R. Huynh, J. Jiang, E. Kremmer, J. Müller, H. Saumweber, A. Spradling and P. Tomancak for providing reagents. We thank S. Baldi, D. Jain, N. Steffen and S. Vengadasalam for stimulating discussions.

## Author contributions

KB performed experiments; KB and PBB developed concepts and approaches; KB and PBB analyzed data; KB and PBB prepared the manuscript.

